# Transcriptomic biomarkers for prediction/classification of sample conditions in marine microeukaryotes (diatoms)

**DOI:** 10.1101/491258

**Authors:** Justin Ashworth

## Abstract

Marine microeukaryotes express large and complex transcriptomes that often respond dynamically to environmental and physiological conditions. In parallel to developments in human disease research, the opportunity exists to employ transcriptomic features as “biomarkers” to understand and predict cellular and environmental states. Here, the prediction and classification of basic physiological and environmental states including light, growth phase and inorganic carbon status was explored for the model diatom *T. pseudonana* using publicly available data including 56 microarray and 316 mRNA-seq samples. Simple “machine learning” methods combined with integrative bootstrapped clustering were able to detect, recapitulate and expand biologically and environmentally relevant signals evident across hundreds of samples collected and processed independently by multiple laboratories. Agnostic, integrative and empirical “data-driven” approaches are likely applicable to modern questions in new environmental and experimental contexts.

## Introduction

Cells experience large changes in their environment, and operate complex evolutionarily successful programs accordingly to navigate these challenges. Marine microbes including diatoms thrive in oceanic environments, where conditions fluctuate and shelter is typically impossible (Armbrust 2009). The use of transcriptomics in environmental biology has focused on the identification of processes and genes that respond to controlled environmental shifts (Mock et al. 2008; Allen et al. 2008; Waldbauer et al. 2012; Shrestha et al. 2012; Hennon et al. 2015; Nymark et al. 2013). This has revealed dozens of intracellular programs and thousands of genes whose correlations with treatment conditions deepen knowledge of the internal biology, homeostasis and coping strategies of marine microbes.

In medical fields, efficient molecular data collection has been further applied to the development of predictive molecular “biomarkers” that can predict and classify samples, cells, tissues and subjects, e.g. as “healthy” or “diseased” (Akbani et al. 2015). The correlation of highly specific molecular readings, including transcript levels, with the biological state of test subjects allows the classification, diagnosis or corroboration of hypothetical or hidden conditions (Li et al. 2017). This may also be applicable to biological and environmental questions, wherein measurable molecular biomarkers can indicate the biological and environmental status of study areas (Saito et al. 2014).

In a “forward transcriptomics” context, deliberate changes in environmental (laboratory) conditions are used to expose and identify specific transcripts whose expression levels are most significantly affected by these controlled dependent variables. These efforts employ powerful and highly refined statistical techniques that are primarily designed for this purpose (Ritchie et al. 2015; Robinson, McCarthy, and Smyth 2010). Conversely, a “reverse transcriptomics” approach would seek to predict and identify uncertain phenotypic and environmental conditions, based on learned information about relationships between transcriptional programs and intra- and extra-cellular environments. A convenient, agnostic and “model-free” approach to this is to apply basic “machine learning” tools to project learned associations onto unlabelled samples. Here we explored this purpose using a large public dataset of laboratory transcriptomic data for the diatom *T. pseudonana (Tp)*.

## Light status, light harvesting programs, and growth constraints

Photosynthetic organisms respond dynamically to light conditions, balancing light harvesting and the shuttling of electrons for energy acquisition against the damaging effects of overexposure (Demmig-Adams and Adams 1992). In diatoms, as in other algae and plants, this is evident in varying levels of photoproteins and photochemistry with respect to light exposure (Oeltjen, Krumbein, and Rhiel 2002) and the active relationship between light flux and photosynthetic rates (MacIntyre et al. 2002).

The *T. pseudonana* draft genome (Armbrust et al. 2004) includes at least 37 putative light harvesting complex (Lhc) proteins, which are dynamically expressed and highly coordinated **(Fig. 1)**. This apparent sensitivity of Lhc transcript levels to environmental status has presumably evolved as part of a system to tightly control light harvesting potential. Shifts in the levels of transcripts encoding Lhcs are among the largest and most significant differences observed in transcriptomic experiments (Ashworth et al. 2013; Nymark et al. 2013).

**Figure 1.**
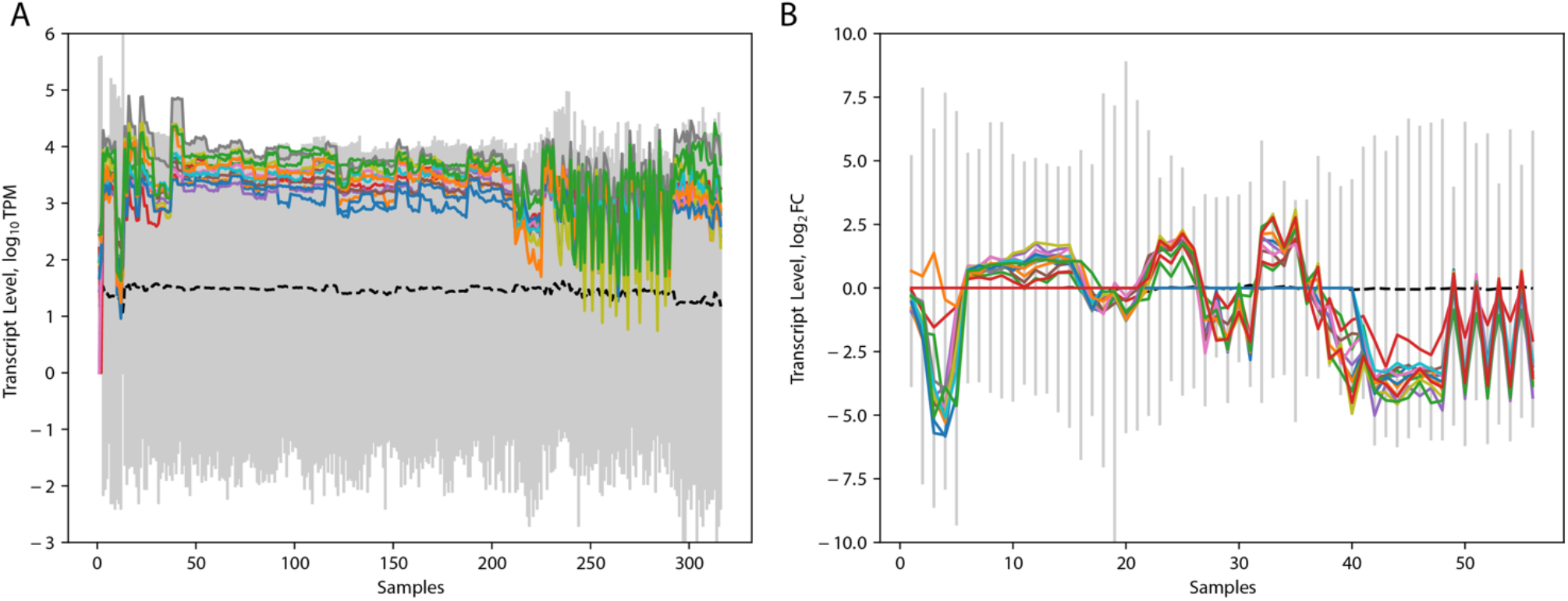
The coordinated expression patterns of transcripts encoding light harvesting complex (Lhc) proteins in the diatom *T. pseudonana*. The clustering of these 14 transcripts according to the similarity of their apparent expression levels over many conditions is highly reproducible (Ashworth et al. 2016; Ashworth and Ralph 2018). A) 316 public mRNA-seq samples, B) 56 public microarray samples. Gray bars indicate the range of minimum and maximum values occurring in each sample, and a dashed line indicates median normalized mRNA-seq transcript levels (A) or microarray expression changes (B) per sample.

### Can transcriptomic features predict light status?

To answer this question, we fit simple linear multi-dimensional kernel models (a “support vector machine” or SVM included in the Python ‘scikit-learn’ package) to 52 samples grown under known and varying light levels. SVMs optimize multi-dimensional support vectors that divide data into groups according to labelled samples, and can also predict the classes of unlabelled samples for new data projected across trained (fit) models. Candidate biomarkers in this analysis included:

i. the full complement of normally-observed *T. pseudonana* transcripts,
ii. individual transcripts, and
iii. bootstrap-supported clusters of transcripts whose similarity of expression was robust to noise simulated by empirical resampling (Ashworth and Ralph 2018; Suzuki and Shimodaira 2006).

The features whose transcript levels most accurately recapitulated light conditions in these samples are shown in **Table 1**. While numerous features are available among the 11,221 mRNA-seq transcripts that were observed in at least half of all samples, only 76 of tested features resulted in models that predicted >90% of samples with known light conditions (“light” or “dark”) correctly. These consisted of 35 single transcripts, 40 bootstrap clusters, and 1 feature consisting of the whole transcriptome. The vast majority were relatively uninformative, with a median cross-validated prediction accuracy of ~40% for single-transcript features **(Fig. 2)**.

**Table 1.** The 20 features whose transcript levels best predicted the light status of 52 mRNA-seq samples with annotated light levels.

| Accuracy<br>(Test Set)<br>correct/all | Feature<br>Size | Tp v.3<br>IDs | Annotation / Function |
| --- | --- | --- | --- |
| 0.961 | 8 | [8 IDs] | threonine synthase activity, transcription cofactor activity |
| **0.959** | **1** | **4202** | **(Tp_bHLH8_PAS) predicted pas-domain protein** |
| 0.957 | 1 | 2892 | EF_TS(3e-134), Mitochondrial translation elongation factor EF-Tsmt, catalyzes nucleotide exchange on EF-Tumt, tsf |
| **0.955** | **14** | **[14 IDs]** | **membrane, photosynthesis (light harvesting)\*** |
| 0.955 | 12 | [12 IDs] | protein-methionine-S-oxide reductase activity, inositol or phosphatidylinositol phosphatase activity, inositol-1(or 4)-monophosphatase activity |
| 0.955 | 1 | 7940 | [Unk.] |
| 0.952 | 1 | 1913 | P-loop NTPase(2e-21), AAA+-type ATPase, AAA |
| 0.952 | 18 | [18 IDs] | carbohydrate metabolism, NAD binding, ribose-5-phosphate isomerase activity, pentose-phosphate shunt, non-oxidative branch, 3-oxoacyl-[acyl-carrier protein] reductase activity, phosphoglycerate kinase activity, phosphoglucomutase activity, ribulose-phosphate 3-epimerase activity, intramolecular transferase activity, phosphotransferases |
| 0.952 | 13 | [13 IDs] | unfolded protein binding, chaperonin ATPase activity, electron transporter activity, electron transport, endoplasmic reticulum, protein disulfide isomerase activity, protein folding |
| 0.950 | 1 | 9689 | Transcription factor CHX10 and related HOX domain proteins, (Tp_bZIP7b/PAS) predicted regulator [Rayko et al.] |
| 0.950 | 1 | 24507 | Nuclear receptor coregulator SMRT/SMRTER, contains Myb-like domains; (Tp_bZIP5_PAS) predicted regulator [Rayko et al.] |
| 0.948 | 10 | [10 IDs] | D-xylose 1-dehydrogenase (NADP+) activity, caspase activity, guanosine tetraphosphate metabolism |
| 0.943 | 18 | [18 IDs] | uroporphyrinogen decarboxylase activity, protochlorophyllide reductase activity, trans-9R,10R-dihydrodiolphenanthrene dehydrogenase activity, glucose-6-phosphate isomerase activity, cis-2,3-dihydrodiol DDT dehydrogenase activity, bacteriochlorophyll biosynthesis, gluconeogenesis, trans-1,2-dihydrodiolphenanthrene dehydrogenase activity, oxidoreductase activity, enoyl-[acyl-carrier protein] reductase activity |
| 0.941 | 9 | [9 IDs] | NAD(P)+ transhydrogenase (AB-specific) activity, phosphomannomutase activity, electron transport, intramolecular transferase activity, phosphotransferases, proton transport, FMN binding, NAD(P) transhydrogenase activity |
| 0.941 | 5 | [5 IDs] | dipeptidyl-peptidase IV activity, prolyl oligopeptidase activity, serine-type peptidase activity |
| 0.941 | 16 | [16 IDs] | ATP synthase activity, lipid biosynthesis, light-harvesting complex, chlorophyll binding, photosystem, chloroplast ribulose biphosphate carboxylase complex, cyclopropane-fatty-acyl-phospholipid synthase activity, carbon utilization by fixation of carbon dioxide, photosystem II stabilization, photosynthesis, light reaction, ribulose-bisphosphate carboxylase activity, electron transporter, photosynthetic electron transport chain, protein stabilization, photosystem II, photosystem I reaction center, photosynthesis |
| 0.939 | 8 | [8 IDs] | mRNA metabolism, aspartate oxidase activity, nucleotide kinase activity, phospholipid biosynthesis, inositol-3-phosphate synthase activity, nucleobase, nucleoside, nucleotide and nucleic acid metabolism, myo-inositol biosynthesis, adenylate kinase activity, L-aspartate oxidase activity |
| 0.939 | 11221 | * | [Entire *T. pseudonana* transcriptome] |
| 0.936 | 1 | 11564 | 3.2.1.3, Chitinase |
| 0.936 | 7 | [7 IDs] | procollagen-proline, 2-oxoglutarate-4-dioxygenase complex, prolyl aminopeptidase activity, aminopeptidase activity, procollagen-proline 4-dioxygenase activity |

**Figure 2.**
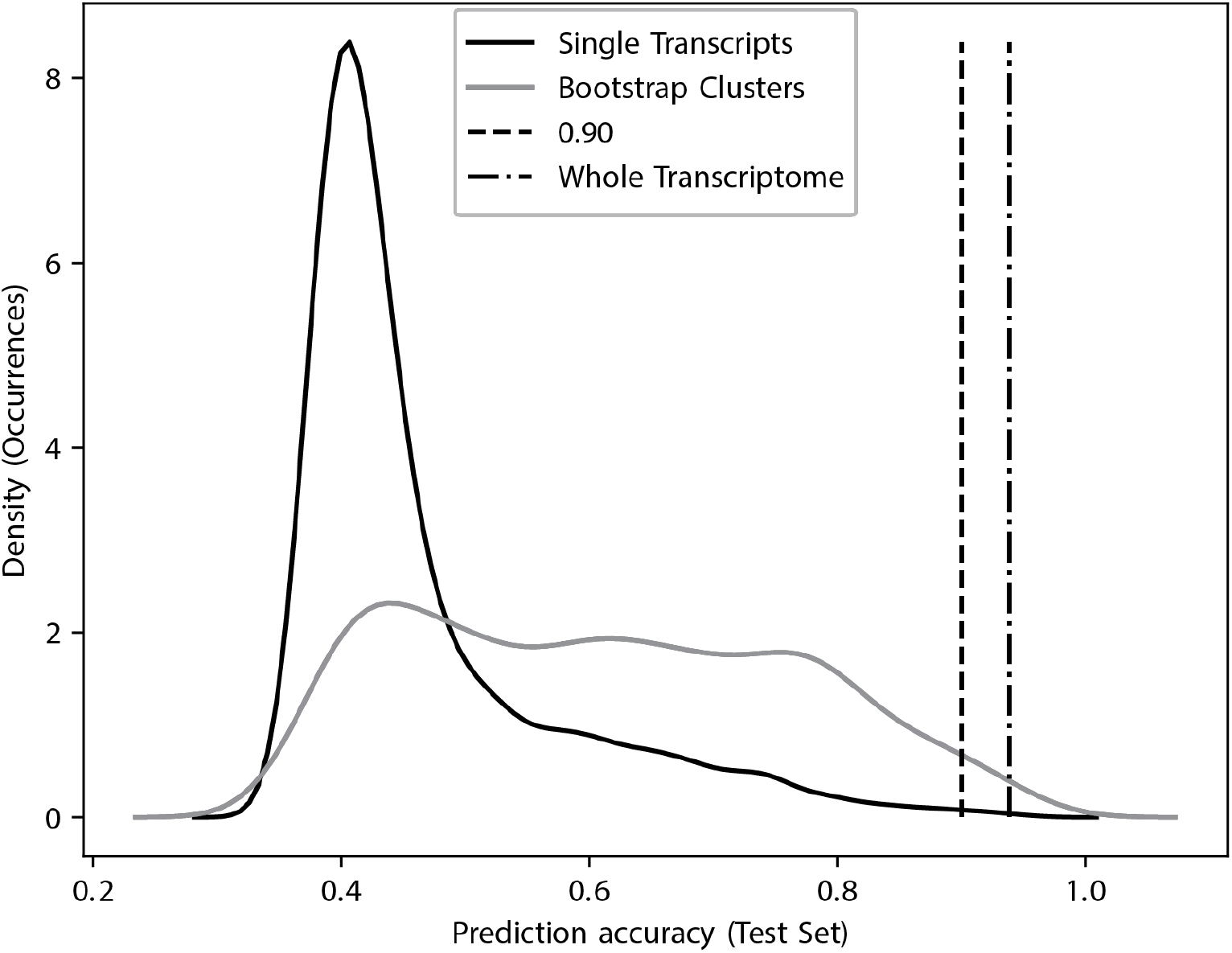
Distributions of cross-validated prediction accuracy for light status on untrained samples for single- and multi-transcript SVM classifiers.

Including 40-fold cross-validation, the information contained in the expression levels of 14 Lhc transcripts **(Table 1)**, ‘<u>photosynthesis (light harvesting)</u>’) correctly classified conditions as “light” or “dark” with 96.7%/95.5% accuracy (trained samples/untrained samples), performing well at predicting the light conditions of untrained samples as a model trained using a matrix of all 11,221 transcripts in this dataset (100%/93.9% accuracy). This finding is consistent with the theory that dynamic regulation of Lhcs balances the collection of photosynthetically active radiation (PAR) by antennae proteins, and is informative of the organism’s cellular state with respect to known (or unknown) light conditions.

The most predictive single transcript feature (*Tp* v.3 4202) encodes a putatively light-sensitive basic helix-turn-helix/PAS transcription factor domain (Rayko et al. 2010). The levels of this transcript increase ~20-fold in cells experiencing “dark” conditions. While this single-transcript feature may be highly predictive of light status across these samples, it is a low abundance transcript, which might be unobserved in new samples with low coverage. Across 316 public mRNA-seq samples for *T. pseudonana*, the median normalized read count (transcripts per million, TPM) for *Tp* 4202 is among the 2.7% least abundant transcripts, and it was not observed at all in 21 samples. The prospect of false negatives due to low transcript levels and/or coverage complicates, but does not preclude prediction/classification **(Fig. 3A)**.

**Figure 3.**
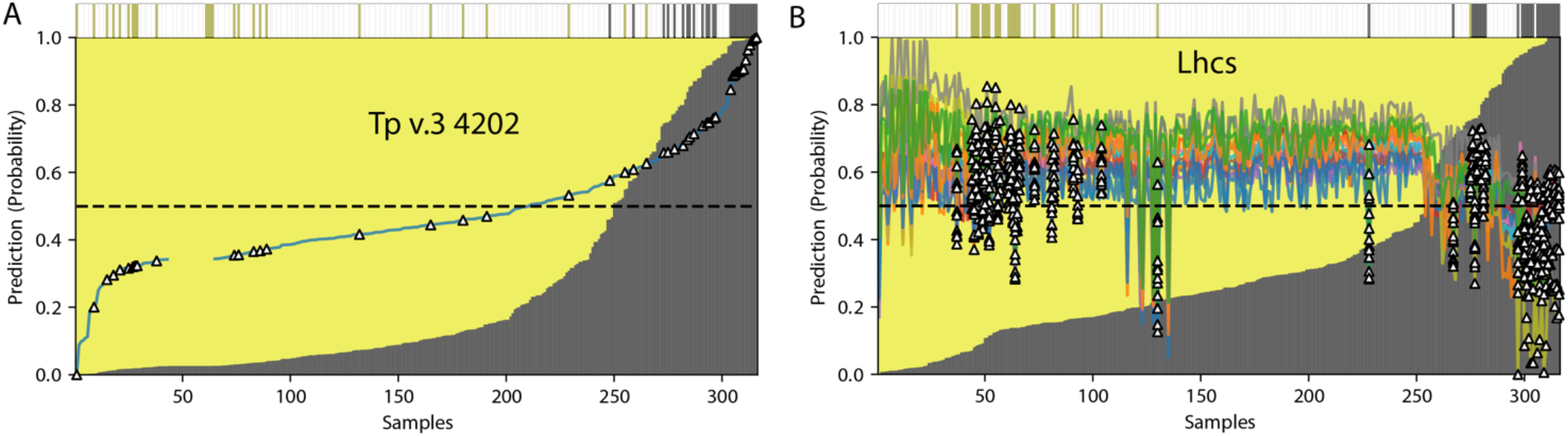
The prediction of light status using trained transcript features across all 316 public mRNA-seq samples. A) The normalized transcript per million (TPM) levels of a single transcript encoding a putative bHLH/PAS light-sensitive transcription factor domain (*Tp* v.3 4202) easily distinguish “light” from “dark” samples. B) A model trained on transcript levels for 14 Lhc transcripts successfully distinguishes between nearly all “light” and “dark” samples. True sample labels are indicated by colored bars at the top. White triangles represent labelled training/testing data, colored lines represent all data, gray regions indicate probabilities of samples being collected under “dark” conditions, yellow regions indicate probabilities of samples being collected under “light” conditions, dashed lines indicate a prediction probability 0.5 (50%).

Informative, coherent clusters transcripts may provide superior predictive biomarkers for light status in this species as compared to single transcripts or whole transcriptomes for multiple reasons:

i. the propensity for overfitting when using many more parameters (all transcripts) than classes (*n*_samples_ ≪ *n*_transcripts_),
ii. the likelihood of encountering missing and/or excessively noisy individual transcripts in additional samples, and
iii. unintended and orthogonal batch, study, inter-lab or inter-sample effects across many (or arbitrary) transcripts in aggregate, to which modern statistical prediction/classification approaches may be extremely sensitive.

The use of data-supported co-expression clusters derived from all available public data incorporates information that is empirically reliable and generalizable over nearly all measured conditions, in contrast to top DE transcripts or clusters drawn solely from individual experimental series. The use of clustering for feature selection and prediction is common in prediction/classification and machine learning problems.

### Predictions for “unlabelled” samples

Applied to the full dataset, including an additional 274 samples not labelled as “light” or “dark” collected across different labs using different platforms, the SVM model fit to 14 Lhcs predicted 78 samples consistent with “dark” conditions, and 238 samples consistent with “light” conditions **(Fig. 3)**. While inaccuracy in these extended predictions is likely to be higher than in internally cross-validated test sets, plausible predictions of unknown condition states based on validated transcriptional patterns may greatly extend the biological and contextual understanding of samples for which conditions are uncertain, unmeasured, or not reported.

### Application to microarray fold-change data

Public microarray data including known changes in light status (Ashworth et al. 2016) may be also amenable to prediction/classification **(Fig 4)**. While information about raw transcript abundance is typically not included or reliable in microarray studies, gene-wise expression fold-change ratios to common references are informative of programmatic shifts in intracellular regulatory and physiological programs. Among 52 publicly available microarray samples for *T. pseudonana* (Ashworth et al. 2016), 35 were collected under known conditions of “light,” “dark,” or “excess” light. While the expression levels of Lhcs were able to classify light status in most samples (87.2%/69.3%; trained/test samples), the effect of growth phase on relative changes in Lhc expression appears to complicate these predictions **(Fig. 4B)**. Samples collected during timeseries experiments exhibited large phase-dependent shifts in Lhc expression (Ashworth et al. 2013). Microarray expression changes for the bHLH/PAS *Tp* v.3 4202 gene were completely uninformative of light status across these experiments, potentially due to loss of signal for this lowly-expressed gene **(Fig. 4A)**.

**Figure 4.**
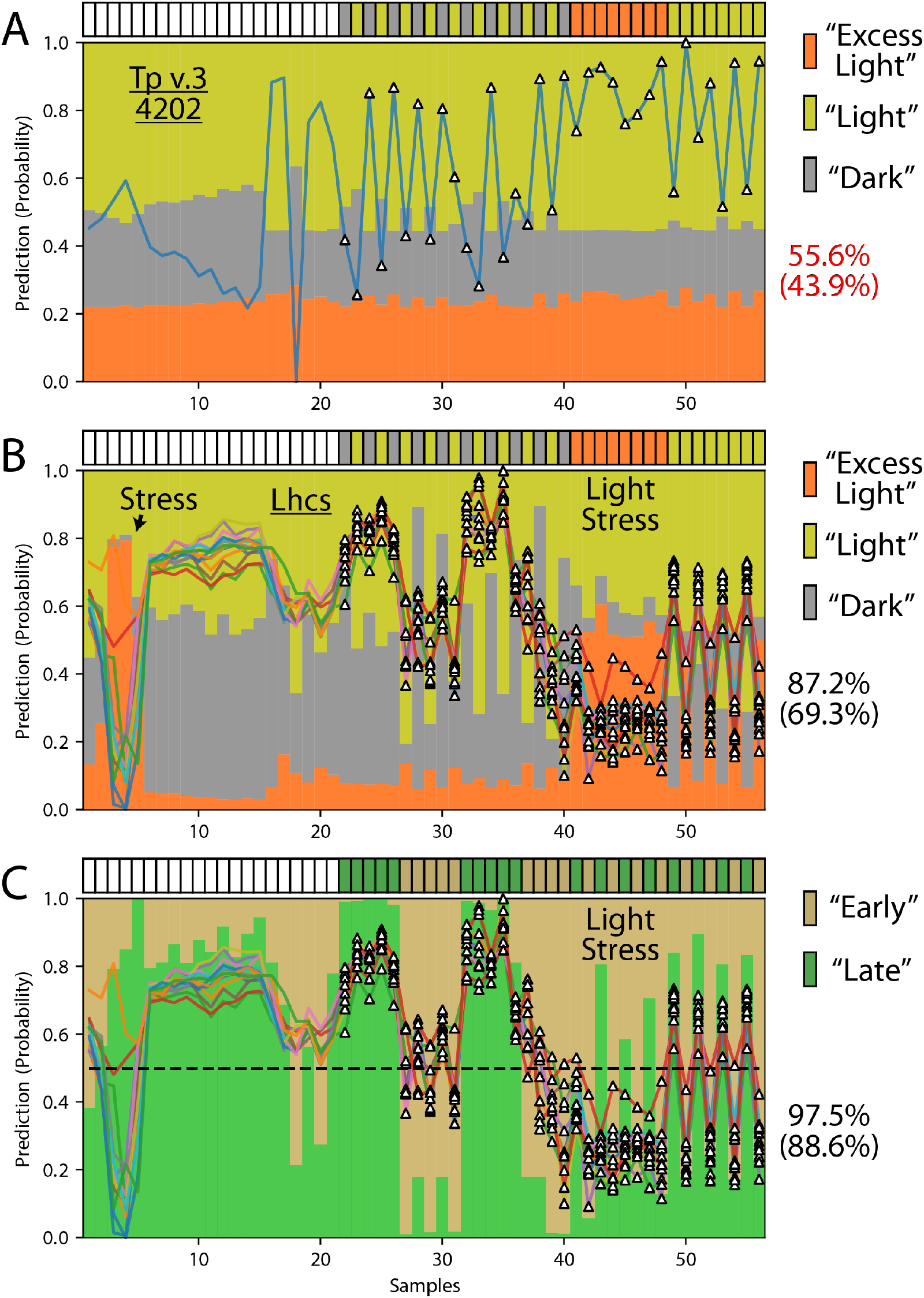
The prediction of light status and growth phase using trained relative gene expression features across all 52 public microarray samples. True sample labels are indicated as colored boxes at the top of each plot. White boxes indicate “unlabelled” samples. Prediction accuracies are shown on the right (training/testing data). A) The *Tp* v.3 ID 4202 (putative bHLH/PAS) gene is not informative of light status in these microarray data, due potentially to its low expression level. B) Relative expression changes for a bootstrap-supported Lhc cluster of 14 genes successfully recapitulates light status in microarrays. Cross-classification to unlabelled samples known to be in stress conditions (“Stress”) is consistent with the results of previous specific stress-related experiments (Mock et al. 2008). White triangles represent labelled training/testing data, colored lines show all data.

Among these microarray samples, Lhc transcript levels predicted growth phase with (97.5%/88.6%) accuracy, as Lhcs increased in expression during “early/exponential” growth and decreased in expression in “stationary/limited” cultures. This relationship between growth phase and relative Lhc expression appears to surpass the role of light levels. A departure from this relationship occurs in samples experiencing high light stress, where Lhc expression is down-regulated during all phases of growth **(Fig. 4C)**.

Lhcs also successfully predicted growth phase in mRNA-seq data, predicting samples labelled as “early” or “late” growth phase with a high accuracy (94.4%/93.6%) comparable to that of the entire transcriptome (100%/92.3%). Out of 316 total samples, 262 samples were predicted to be in a “growing” state and 54 were predicted to be “stationary” based upon the expression levels of light harvesting complexes **(Fig. 5)**. While thousands of changes occur inside diatoms during growth phase transitions (Ashworth et al. 2013; Valenzuela et al. 2018), the Lhcs may be convenient and effective “biomarkers” for the light and growth status of diatoms in uncertain samples or environments.

**Figure 5.**
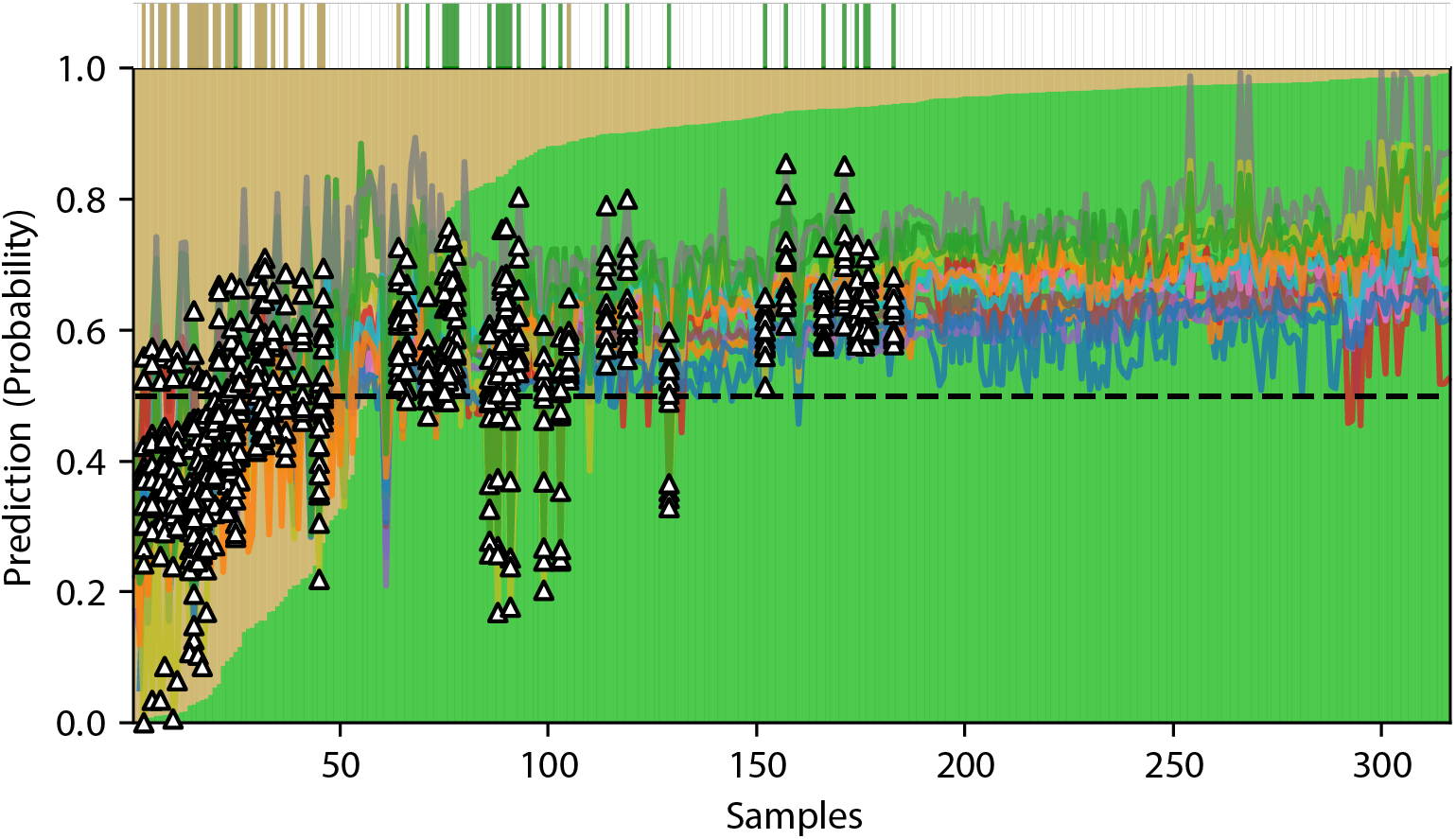
The prediction of growth phase (green: early/growing/exponential; brown: late/limited/stationary) using a bootstrap cluster of transcripts encoding Lhcs as a predictive feature across 316 public mRNA-seq samples. True sample labels are indicated as colored boxes at the top of each plot. White/missing boxes indicate “unlabelled” samples. White triangles represent labelled training/testing data, colored lines show normalized TPM data.

## Inorganic carbon status

Photosynthesis requires CO_2_ as a source of carbon atoms that can be reduced into higher-energy organic molecules. In marine microalgae CO_2_ must be acquired in dissolved forms from the wet environment, and it is actively pumped into the cell across multiple membranes (Hopkinson et al. 2011; Hopkinson, Dupont, and Matsuda 2016; Shen, Dupont, and Hopkinson 2017). When experiencing dissolved inorganic carbon (DIC) levels below ~300-400 ppm CO_2_, diatoms including *T. pseudonana* up-regulate carbon-concentration mechanisms (CCMs) in order to scavenge dissolved inorganic carbon. This situation should occur commonly in natural environments, blooms, and laboratory cultures, whenever photosynthesis outpaces DIC availability.

The transcript levels of several CCM-related genes in *T. pseudonana* have been found to correlate strongly with the presence or absence of carbon limiting conditions (Hennon et al. 2015; Valenzuela et al. 2018). These transcripts in turn may be excellent biomarkers for the prediction of carbon status in samples collected from conditions for which the inorganic carbon status not known, measured or reported.

The performance of several CCM-related transcripts as predictive biomarkers across 149 public mRNAseq samples (70 “low”/79 “high”) with known/labelled CO_2_ conditions is shown in **Table 2**. Predictive models fit on an eight-transcript CCM cluster identified in controlled chemostat experiments (Hennon et al. 2015) and confirmed in large-scale batch cultures (Valenzuela et al. 2018) are able to recapitulate the inorganic carbon status across these samples with 100% accuracy. Close behind are models based on single-or multi-transcript features including the most consistently up-or down-regulated transcripts across controlled experiments, including:

i. a carbonic anhydrase (*Tp* v.3 233),
ii. a lowly-expressed putative permease (*Tp* v.3 262258) that strongly increased in expression under “high” CO_2_ conditions (800 ppm), and
iii. a pair of transcripts putatively encoding bestrophin domains (*Tp* v.3 4819, 4820).

**Table 2.** Cross-validated accuracy of predictions for inorganic carbon status (“low”, “high”) for selected transcript features.

| Accuracy (Test Set) correct/all | Median log10 Fold Change (TPM) “high” vs. “low” CO<sub>2</sub> | Feature | Size | IDs (*Tp* v.3) |
| --- | --- | --- | --- | --- |
| 1.000 | -0.4 | “CCM” Cluster | 8 | 233, 264181, 4819, 4820, 269696, 269699, 10360, 6529 |
| 1.000 | -0.4 | “CCM” Cluster & Permease | 9 | 233, 264181, 4819, 4820, 269696, 269699, 10360, 6529, 262258 |
| 0.992 | 0 | All transcripts | 11221 | [...] |
| 0.959 | 0.3 | Permease & CdCA1 | 2 | 262258, 233 |
| 0.946 | 1.3 | Permease | 1 | 262258 |
| 0.91 | -0.7 | CdCA1 | 1 | 233 |
| 0.809 | -0.4 | Bestrophins | 2 | 4819, 4820 |

Treated as “biomarkers,” these transcripts result in the prediction of between 148 to 216 “low” (≤ ~400 ppm) and 100 to 166 “high” (≥ ~400 ppm) samples. The most accurate cluster-based predictor (“CCM Cluster”) predicts 176 “low” and 140 “high” CO_2_ samples; the most conservative predictor (Permease + CdCA1) predicts 216 “low” and 100 “high” CO_2_ samples. These results are illustrated in **Figure 6**.

**Figure 6.**
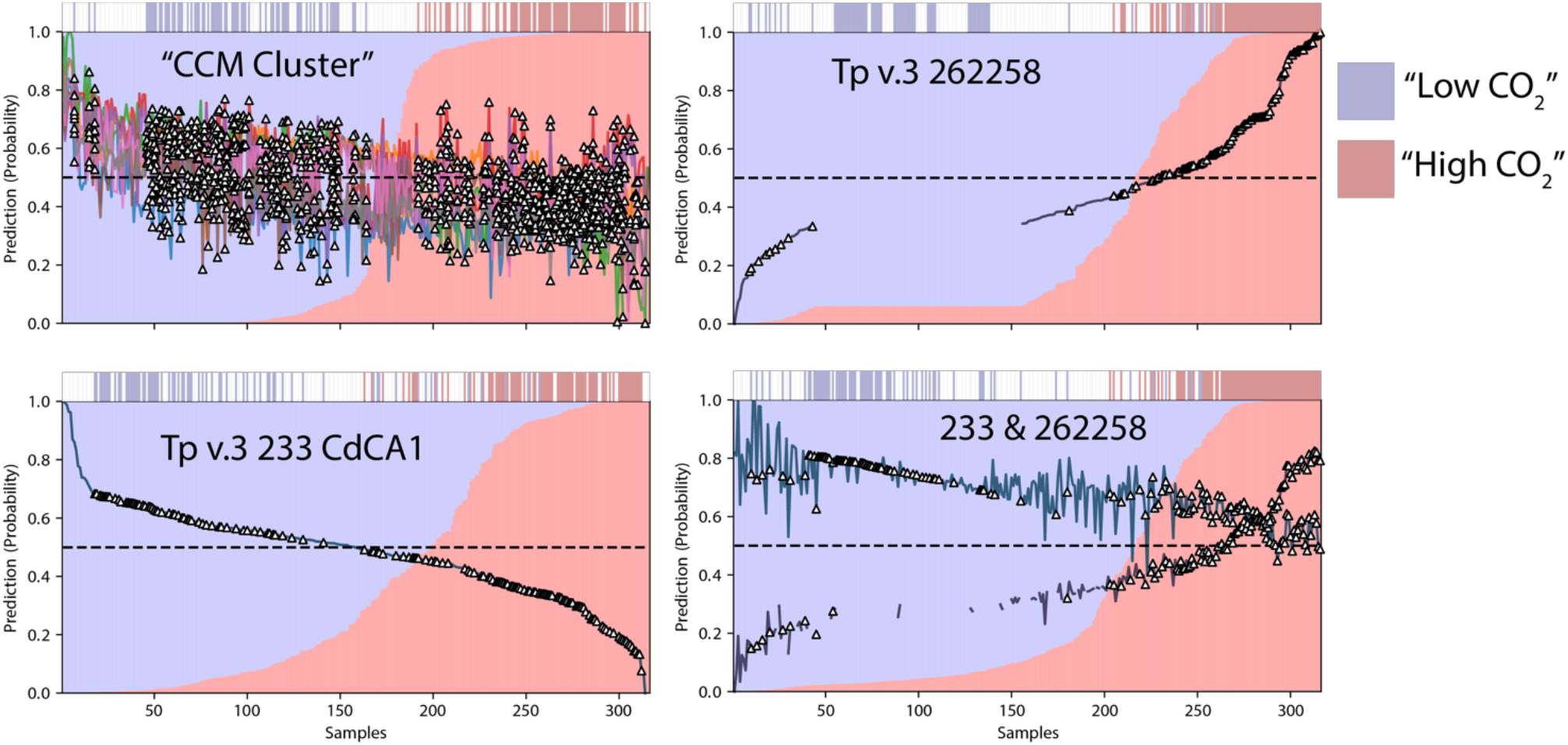
The prediction of inorganic carbon status (blue: “low” (≤ ~400ppm CO2), red: “high” (≥ ~400ppm CO_2_) using a cluster of transcripts encoding CCM-related functions (“CCM Cluster”), a carbonic anhydrase (*Tp* v.3 233), a putative permease (*Tp* v.3 262258) and a combination of 233 & 262258. True sample labels are indicated as colored bars at the top of each plot. White/missing boxes indicate “unlabelled” samples. White triangles represent labelled training/testing data, colored lines show normalized TPM data.

While the putative permease *Tp* v.3 262258 is a highly informative biomarker for inorganic carbon status in this dataset, relative counts for this transcript are exceedingly low (or missing) in a large number of samples (122/316). Even when this transcript is observed, its median normalized TPM was among the lowest 4% of transcripts. While the absence of reads mapping to 262258 may be indicative of samples that are not experiencing higher than normal CO_2_ conditions, the use of this transcript as biomarker may be challenging due to the likelihood of false negatives in coverage-limited experiments.

The carbonic anhydrase CdCA1 (*Tp* v.3 233), while only slightly less predictive of CO_2_ status, in contrast was expressed in the 79th percentile among all transcripts. Reads mapping to CdCA1 should be more reliably detected in new or low-coverage samples. A predictor based on the combination of these two candidate biomarkers outperformed either transcript alone (**Table 2, Fig. 6**, “233 & 262258”}).

## Nutrient status

Diatoms also employ mechanisms to scavenge nutrients under conditions of depletion or scarcity, and are transcriptionally responsive to these conditions (Mock et al. 2008). The two principal known assimilation genes for silicate (SiO_4_) and nitrate (NO_3_) in *T. pseudonana* are SIT1 (*Tp* v.3 268895) and NRT1 (*Tp* v.3 27414). Predictive features based on the transcript levels of these genes, as well as combinations of gene pairs and corresponding bootstrap clusters are able to predict nutrient status (-Si, -N, replete) with high accuracy in a set of 42 training/testing samples for which nutrient status was annotated or fairly certain **(Table 3)**. Combinations of SIT1 and NRT1 clusters performed more robustly than arbitrary features for the prediction of nutrient status, although multiple apparent cell signaling clusters that were able to separate these classes may also deserve further investigation **(Fig. 7)**.

**Table 3.** Cross-validated accuracy of predictions for nutrient status (-Si, -N, replete) for selected transcript features.

| Accuracy (Test Set) correct/all | Size | IDs (Tp v.3) | Function |
| --- | --- | --- | --- |
| 1 | 9 | 23657, 24060, 25299, 269274, 27414 (NRT1), 3393, 268895 (SIT1), 14322, 9619 | Silicate and nitrate assimilation |
| 0.994 | 11221 | [Whole transcriptome] |  |
| 0.994 | 2 | 268895 (SIT1), 27414 (NRT1) | Silicate and nitrate assimilation |
| 0.947 | 9 | 15976, 2247, 26403, 268080, 33699, 36213, 8259, bd381, bd763 | Signalling |
| 0.944 | 10 | [...] | Signalling |
| 0.944 | 7 | 20569, 269557, 33407, 34738, 3517, 37054, 8675 | Signalling |
| ... | ... | ... |  |
| 0.911 | 3 | 268895 (SIT1), 14322, 9619 | Silicate assimilation |
| ... | ... | ... |  |
| 0.906 | 1 | 268895 (SIT1) | Silicate assimilation |
| ... | ... | ... |  |
| 0.85 | 1 | 27414 (NRT1) | Nitrate assimilation |

**Figure 7.**
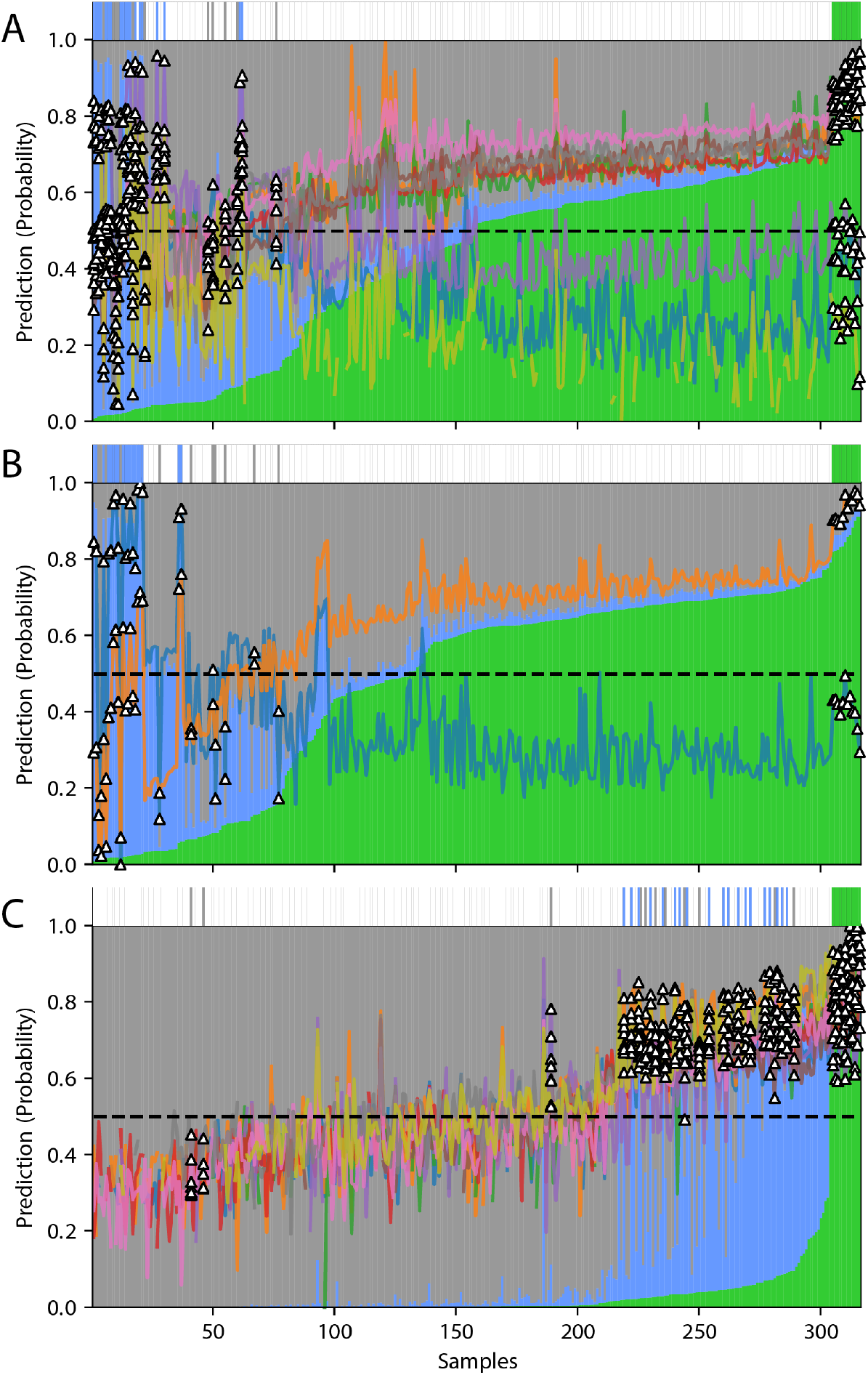
The prediction of nutrient status (blue: -Si, green: -N, gray: replete) using transcript features including A) combined SIT1- and NRT1-containing clusters, B) SIT1 and NRT1 in concert, C) an apparent signalling cluster of relatively unknown significance with high discriminatory ability. Known sample labels are indicated as colored bars at the top of each plot. White/missing boxes indicate “unlabelled” samples. White triangles represent labelled training/testing data, colored lines show normalized TPM data.

## The real world? Limited references

Modern efforts in biological oceanography include the application of genomics and transcriptomics to understand what is occurring at wild and diverse study sites (Marchetti et al. 2012). Outside of the laboratory and in contexts wherein cellular material and coverage are scarce, limitations on biomass or sequencing depth often preclude full or adequate coverage of species’ entire transcriptomes. Methods of inference may fail outside of controlled laboratory conditions due to new environments, data sparsity, and missing/non-uniform references or internal standards.

To explore the issue of limited, missing or sparse reference transcripts, we re-fit models to data normalized to a limited but adequate array of ad hoc “stable” transcripts evident in all laboratory samples. These transcripts are evidently maintained at consistent levels across a range of conditional variation represented in the dozens of experiments, and could be considered “housekeeping genes” for the purposes of agnostic normalization (Glusman et al. 2013). The 20 least variant transcripts whose normalized read counts (TPM) were above the median across 316 samples are shown in **Figure 8** and **Table 4**.

**Figure 8.**
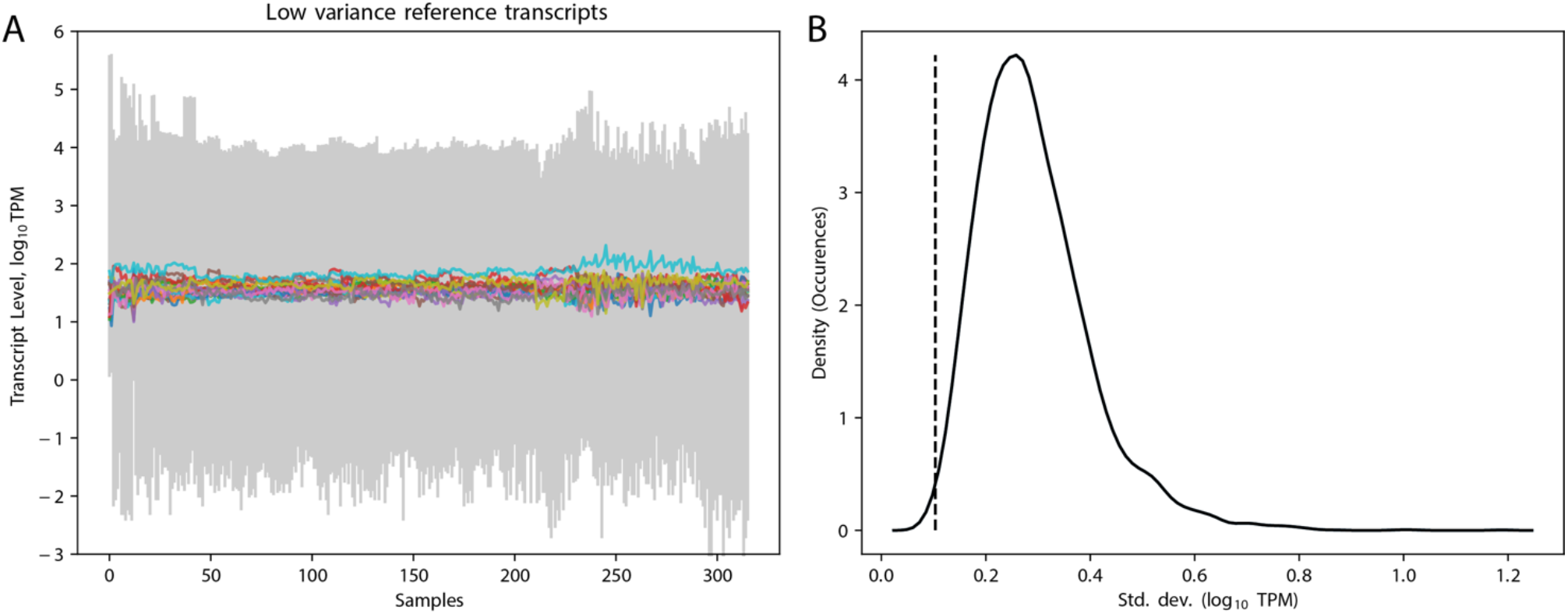
Stable reference transcripts across many laboratory experiments. A) Lines indicate normalized transcript read count values (TPM) for each of the transcripts listed in Table {RefTx}. The gray region indicates the range of minimum and maximum values occurring in each sample.

**Table 4.**
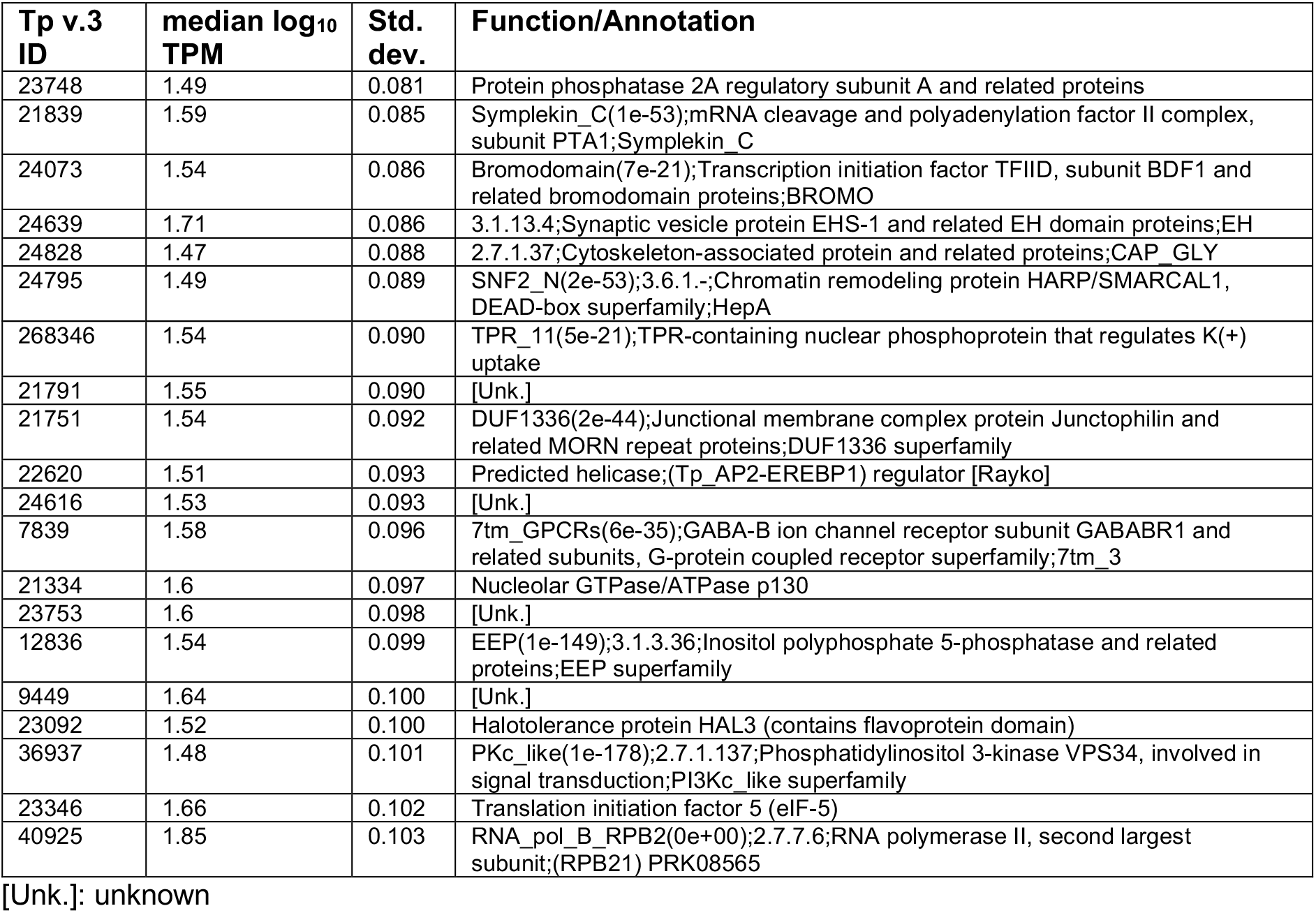
Identities of the 20 most stable reference transcripts with above-average median transcript levels in this dataset.

As ratios to median values of empirically-derived reference transcripts, transcript levels of Lhcs continue to correctly predict “low” and “high” light conditions, as well as “exponential” and “stationary” growth phases, and CCM transcripts continue to predict inorganic carbon status **(Table 5)**. More challenging normalization problems may draw upon various combinations of these or dozens of other apparent reference candidates according to their representation in new samples.

**Table 5.** Re-classification of sample conditions based on ratios of biomarker transcripts to empirically-derived stable reference transcripts.

**Prediction: light conditions**
| Accuracy (Test Set)<br>correct/all | Feature | Size | Tp v.3 IDs |
| --- | --- | --- | --- |
| 0.968 | Carbohydrate metabolism | 18 | [...] |
| 0.945 | Unknown function | 8 | 1913, 20777, 21311, 24507, 24613, 268070, 2714, 4632 |
| 0.941 | bHLH/PAS | 1 | 4202 |
| 0.939 | Light harvesting | 14 | [...] |
| 0.939 | Whole transcriptome | 11221 | [...] |

**Prediction: growth phase**
| Accuracy (Test Set)<br>correct/all | Feature | Size | Tp v.3 IDs |
| --- | --- | --- | --- |
| 0.939 | Light harvesting | 14 | [...] |
| 0.930 | Whole transcriptome | 11221 | [...] |

**Prediction: inorganic carbon conditions**
| Accuracy (Test Set)<br>correct/all | Feature | Size | Tp v.3 IDs |
| --- | --- | --- | --- |
| 1.0 | CCM | 8 | 233, 264181, 4819, 4820, 269696, 269699, 10360, 6529 |
| 1.0 | CCM & permease | 9 | 233, 264181, 4819, 4820, 269696, 269699, 10360, 6529, 262258 |
| 0.992 | Whole transcriptome | 11221 | [...] |
| 0.945 | permease | 1 | 262258 |
| 0.944 | Permease & CdCA1 | 2 | 262258, 233 |
| 0.912 | CdCA1 | 1 | 233 |

## Conclusion

Transcriptomic biomarkers that can be trained and validated using large and sufficiently labelled data/sample sets may be useful for predicting the conditions of samples for which various parameters, environmental conditions or cellular states may be unknown or unmeasured. The elementary “machine learning” methods employed here are easily adaptable to additional questions, and can likely be refined or improved to suit more specific needs or constraints. This may be useful for environmental studies of marine microbes using transcript data, and meta-studies that seek to predict and derive new information from large public datasets. Here we explored the basic utility of this approach using public data from a single diatom species, *T. pseudonana*. Extension to additional conditions, species or cross-species questions is likely also feasible.

Caveats to this approach include the need for large numbers of labelled samples, and the propensity for purely “data-driven” models to be overfit to spurious or unreproducible signals. The added problems of non-uniform losses in read coverage, inadequate data to reliably calculate normalized read count values (e.g. TPM), and inabilities to adequately control for compositional biases of mixed populations of cells/species over space and time are outside of the scope of this analysis, but will no doubt complicate efforts to perform predictions for wild samples.

## Data and Methods

The *T. pseudonana* mRNA-seq and microarray datasets used in this analysis consisted of publicly available data downloaded from the NCBI GEO (Barrett et al. 2013) and SRA (Leinonen, Sugawara, and Shumway 2011) databases as of mid-2018. Data integration, normalization and clustering are described in (Ashworth et al. 2016; Ashworth and Ralph 2018). Prediction/classification was performed in Python (version 3.7.0; https://www.python.org/) using linear kernel support vector machine (SVM) tools from the scikit-learn package (version 0.20.1; https://scikit-learn.org). Reported “prediction accuracies (test set)” are the result of cross-validation wherein models were fit 40 times to randomly split training and testing samples (70%/30%); the correct recovery of untrained “test” samples was averaged over all untrained subsets. Figures were created using the Python matplotlib package (version 3.0.2; https://matplotlib.org/).

## Acknowledgements

J.A. is the recipient of an Australian Research Council Discovery Early Career Award (DE160100615) funded by the Australian Government.

